# Therapeutic signature mapping of paired direct and indirect LPS injury in an ex vivo human lung perfusion platform reveals injury-specific druggable programs

**DOI:** 10.64898/2026.08.03.739838

**Authors:** Ahad A. Abdalla, David A. Dorward, Antonella Pellicoro, Tom M. Quinn, Stuart Dickson, Adam Marshall, Annya M Bruce, John J. Cole, Keith Finlayson, Richard A. O’Connor, Christopher Haslett, Manu Shankar-Hari, Kevin Dhaliwal

## Abstract

Acute Respiratory Distress Syndrome (ARDS) remains highly morbid and lacks approved disease-modifying pharmacotherapies. Direct (pulmonary) and indirect (extrapulmonary) insults may initiate biologically distinct early injury programs, but human tissue-level evidence from the first hours is scarce. Here we establish a paired, acellular ex vivo lung perfusion (EVLP) platform using human donor lungs unsuitable for transplantation to model direct (endobronchial) and indirect (perfusate) lipopolysaccharide (LPS) injury within the same donor. We profiled lung tissue proteomes at 4 h post-insult and performed therapeutic nomination by querying proteomics-derived injury signatures against the CLUE L1000 perturbational compendium with independent cross-platform validation. Both models developed histological injury and robust cytokine release. Direct injury preferentially enriched neutrophil degranulation, extracellular matrix remodelling and metabolic reprogramming modules, whereas indirect injury showed prominent complement/coagulation perturbation with greater endothelial activation markers in perfusate. Cross-platform prioritisation converged on tractable signalling and epigenetic axes, including JAK/STAT, PI3K/AKT/mTOR, SYK, CDK and HDAC inhibitor classes - yielding a tiered shortlist for EVLP intervention testing. This intact human lung perturbation platform enables injury-stratified mechanistic inference and therapeutic prioritisation in early lung injury relevant to ARDS.

## Introduction

ARDS remains an unresolved challenge in critical care medicine and is characterised by rapid-onset respiratory failure, diffuse alveolar damage and refractory hypoxaemia, with high mortality and substantial long-term morbidity among survivors (1–2). Despite improvements in supportive care, there are no approved disease-modifying pharmacotherapies for ARDS, in part because ARDS is biologically heterogeneous and commonly treated as a single entity in clinical trials (3–4).

A clinically and mechanistically relevant distinction is between direct (pulmonary) and indirect (extrapulmonary) ARDS. Direct ARDS arises from primary lung injury (e.g., pneumonia, aspiration, contusion), whereas indirect ARDS results from systemic conditions (e.g., sepsis, trauma, pancreatitis) where lung injury follows systemic inflammation (5). These categories differ in lung mechanics and inflammatory profiles (6–7) and human biomarker studies demonstrate higher epithelial injury markers in direct ARDS and higher endothelial injury markers in indirect ARDS (5). Preclinical models similarly show differential effects of candidate therapies depending on injury route and aetiology (8–10).

At the tissue level, ARDS reflects dysregulated inflammation and barrier failure at the alveolar-capillary membrane, involving endothelium, epithelium and the basement membrane, leading to protein-rich oedema, impaired gas exchange and activation of coagulation pathways (11–14). Proteomics offers an unbiased route to defining mechanistic programs, but human ARDS proteomics has largely relied on bronchoalveolar lavage (BAL) fluid and plasma because lung biopsy is rarely feasible in critically ill patients (15–20). These compartments provide valuable information but cannot capture the response of lung-resident cells (21–22). Human tissue-level studies of early injury are particularly important given the limitations of animal models and the likelihood that therapeutic windows occur early in the injury cascade (23). Despite this early ARDS mechanisms in human lung tissue remain poorly defined because tissue sampling in the first hours of injury is rarely feasible.

We hypothesised that an intact human lung injury model would resolve divergent early tissue programs in direct versus indirect injury and enable injury-stratified therapeutic prioritisation. EVLP can model both direct and indirect injury through route-specific LPS administration in human lungs (25–26). Ex vivo lung perfusion (EVLP) provides a translational platform to model human lung injury under controlled conditions, enabling longitudinal sampling and compartment-specific analysis (24). Proteomic analyses in EVLP have been demonstrated in animal models and organ preservation settings (27–28) and LPS provides a relevant trigger of innate immunity via TLR4 signalling (29–31). Here, we establish a paired human EVLP model of direct and indirect LPS injury and used proteomics to define early injury programs. We then performed signature-based therapeutic discovery using the CLUE L1000 perturbational compendium, nominating candidate compounds and target classes predicted to reverse direct- and indirect-injury signatures.

## Results

### Establishment of a paired human EVLP model of direct and indirect lung injury

We established an EVLP platform to model early direct and indirect lung injury (Fig. 1A). Ten donors contributed sixteen single lungs (clinical details in Supplementary Table 1). The protocol enabled paired experimentation such that direct and indirect models could be performed using lungs from the same donor (Fig. 1B). Discovery proteomics was performed in a paired subset (n = 5 donors) enabling within-donor comparisons. Histology and perfusate mediator analyses were performed across the broader experimental set where available.

**Figure 1:**
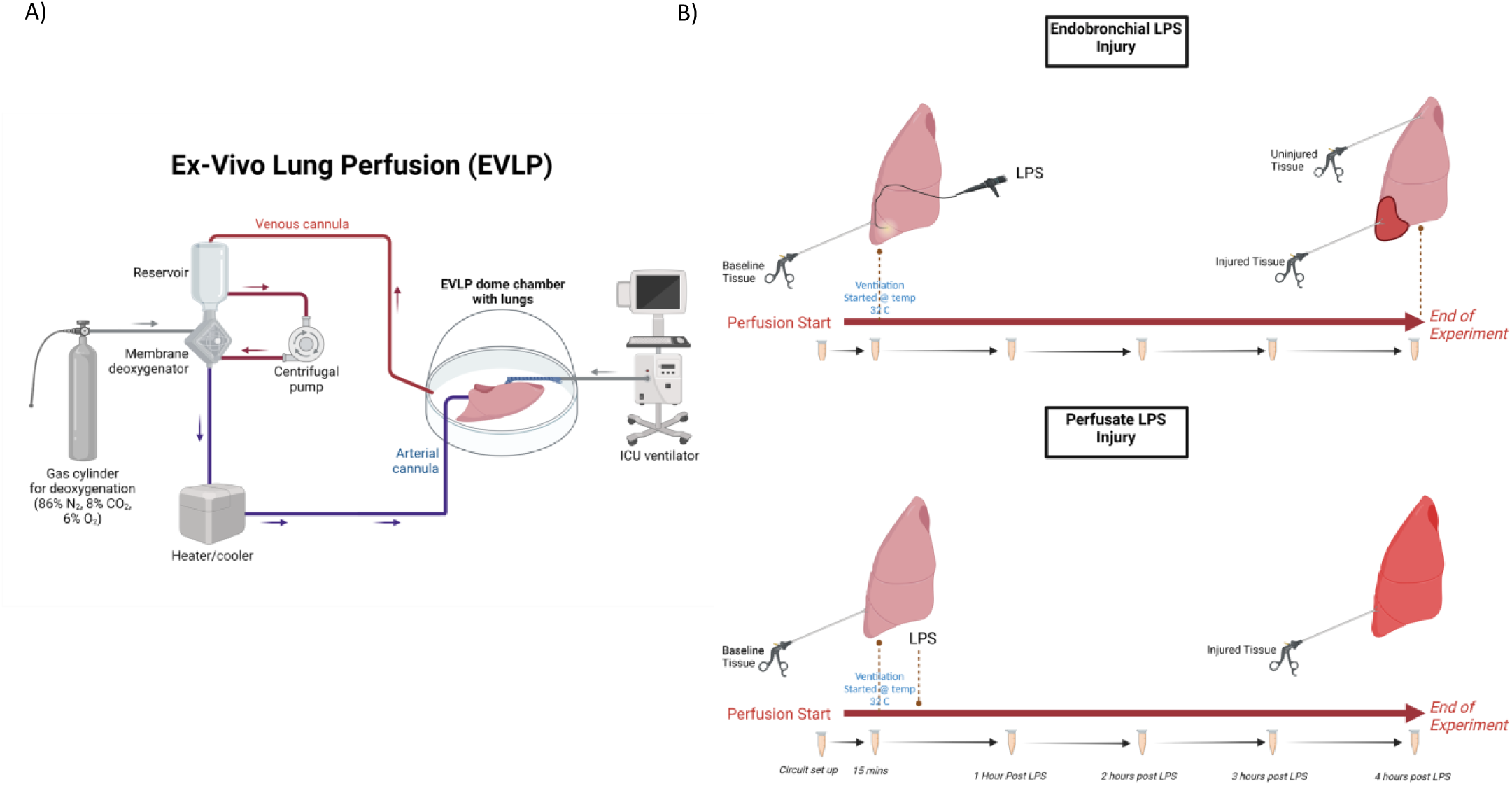
Experimental set-up: A) Schematic representation of the Edinburgh EVLP model. B) Experimental procedure in direct injury model (top) and indirect injury model (bottom).

### Distinct pathological features and mediator trajectories in direct and indirect injury

Baseline histology showed preserved alveolar architecture without inflammatory exudate. At 4 h post-LPS, both models developed lung injury with distinct morphological patterns. Direct endobronchial LPS produced prominent alveolar injury with cellular infiltration, alveolar wall thickening and protein-rich oedema (Fig. 2A). Indirect perfusate LPS resulted in a more diffuse pattern with comparatively less pronounced alveolar-capillary membrane destruction (Fig. 2B). Lung injury scoring increased significantly from baseline to injury in both models (p < 0.01; Fig. 2C-D) using standardised criteria (32). MPO-positive cell quantification demonstrated significant increases in neutrophil infiltration from baseline in both models (p < 0.05).

**Figure 2.**
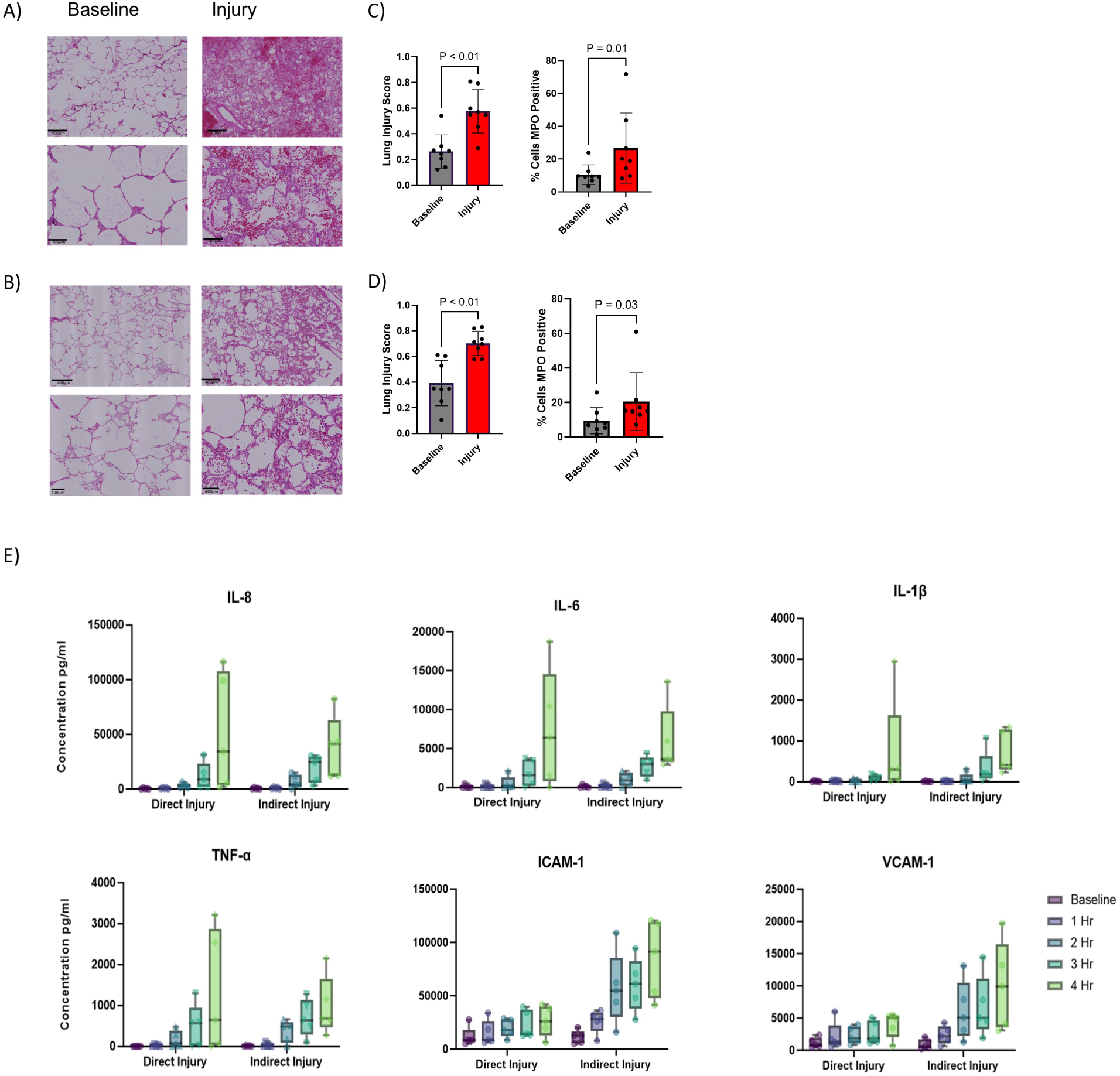
Injury Model: A) Representative H&E-stained sections from direct injury model, lower panels showing magnified regions. B) Representative H&E-stained sections from indirect injury model, lower panels showing magnified regions. C) Lung injury score for direct injury and the proportion of neutrophils (MPO+) from total cell counts for each experiment group. D) Lung injury score for indirect injury and the proportion of neutrophils (MPO+) from total cell counts for each experiment group E) Selected soluble proteins profiles in the perfusate of EVLP experiments, y-axis is concentration in pg/ml and x-axis is time point of measurement. Boxplots show interquartile range with medians. Statistics: Results from paired t-test.

Perfusate profiling showed progressive increases in IL-8, IL-6, IL-1beta and TNFalpha over time in both models, with marked elevation by 4 h (Fig. 2E). Notably, soluble endothelial activation markers were higher in indirect injury: ICAM-1 and VCAM-1 increased substantially more by 4 h in the indirect model, consistent with enhanced endothelial activation in systemic/endotoxaemic-type injury (Fig. 2E).

### Global tissue proteomics resolves EVLP effects and injury-type-specific programs

Across proteomics, 5,650 proteins passed bioinformatic filtering and were quantifiable across planned contrasts. Principal component analysis showed separation of injury states from non-injury controls (baseline and uninjured segments), with principal component 1 capturing approximately 25% of variance (Fig. 3A). Differential testing across three pre-specified contrasts (EVLP effect: baseline vs uninjured; direct injury: injured vs uninjured segment; indirect injury: injured vs matched uninjured segment) identified 874 proteins that were differentially expressed (Fig. 3B) using MS-DAP (33). Significance was FDR-adjusted q < 0.01 together with a per-contrast fold-change threshold estimated by MS-DAP’s bootstrap procedure: |log2FC| > 0.307 (EVLP effect), 0.341 (direct) and 0.347 (indirect).

**Figure 3:**
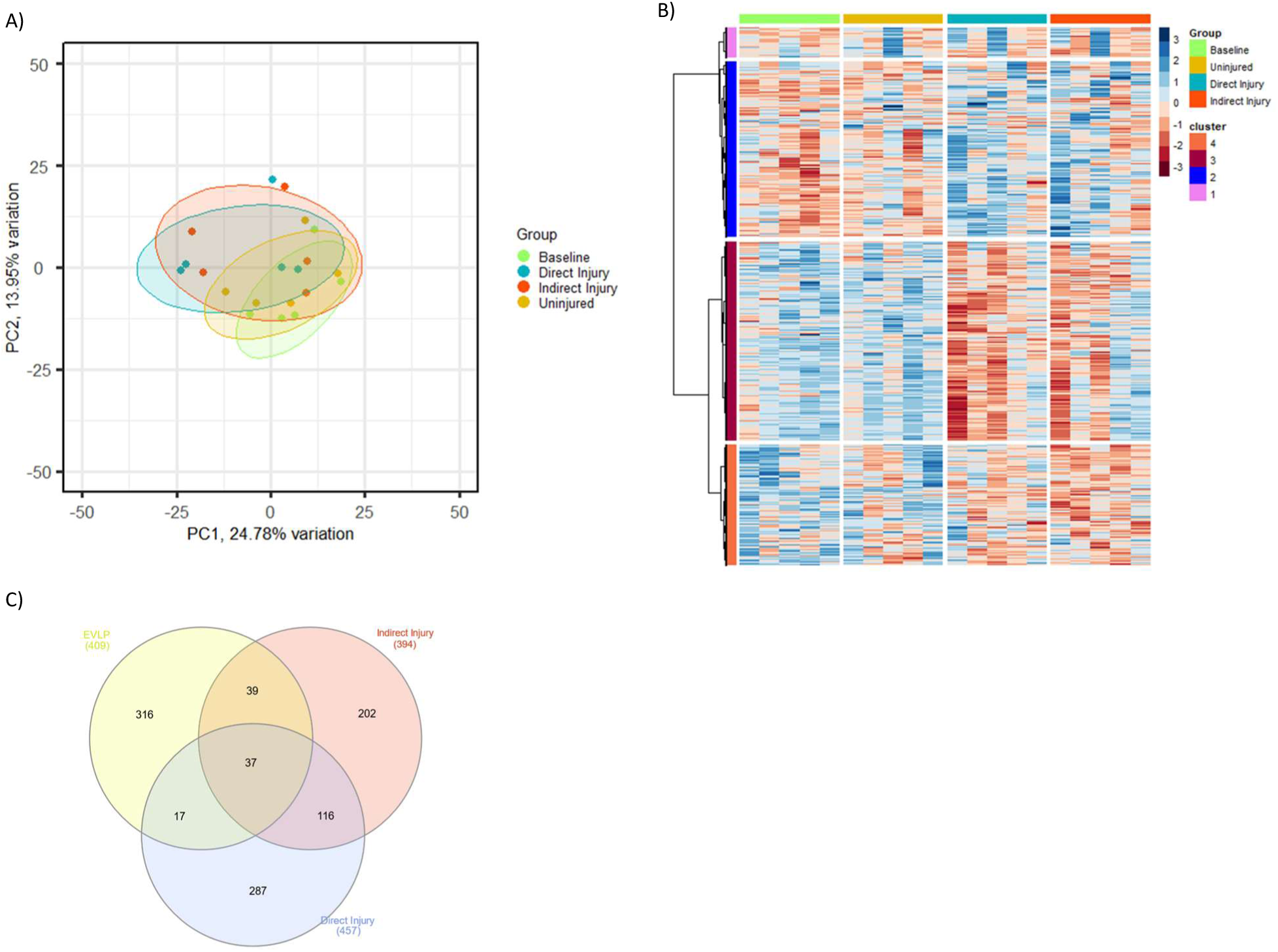
Overall proteomics results: A) Principal component analysis of the4 experimental groups and their protein expression profiles. B) Heatmap of row scaled protein expression values of differentially expressed proteins in any comparison (EVLP, Direct injury and Indirect injury), columns represent samples grouped by sample group and rows represent individual proteins. Functional enrichment revealed that Cluster 1 was mainly involved in proteolysis, Cluster 2 was enriched for innate immune system functions, Cluster 3 was involved in fatty acid and amino acid metabolism and Clust4 ewars a complement and coagulation cascade cluster. C) Venn diagram showing the intersection of the differentially found proteins between the comparisons

Clustering and enrichment analyses from row z-scored abundances in pheatmap, identified modules partitioned into four clusters linked to proteolysis, innate immunity, fatty acid/amino acid metabolism, and complement/coagulation-associated biology (Fig. 3B). After excluding EVLP-associated changes, 116 proteins were shared between direct and indirect injury and enriched for neutrophil degranulation, while each injury type retained distinct pathway enrichment (Fig. 3C).

### Direct lung injury programs

#### Direct injury is dominated by neutrophil granule biology with matrix and metabolic remodelling

In direct injury, differential testing identified 457 proteins differing between injured and uninjured segments at end-EVLP. Of these, 340 were differentially expressed (87 increased; 253 decreased) (Fig. 4A) and 117 were differentially detected. Functional enrichment using clusterProfiler and GO highlighted vesicle/granule cellular components (Fig. 4B). Canonical pathway analysis using the Molecular Signatures Database, leveraging C2:CP canonical pathways (Reactome and KEGG-legacy) sub collections, identified neutrophil degranulation as the dominant signature alongside extracellular matrix organisation and organic acid (including fatty acid) metabolism programs (Fig. 4C). Key neutrophil granule proteins (including azurocidin, neutrophil elastase and myeloperoxidase) were increased in direct injury tissue (Fig. 4A). Statistical significance was defined using a Benjamini–Hochberg false discovery rate (FDR) adjusted *p*-value < 0.05.

**Figure 4:**
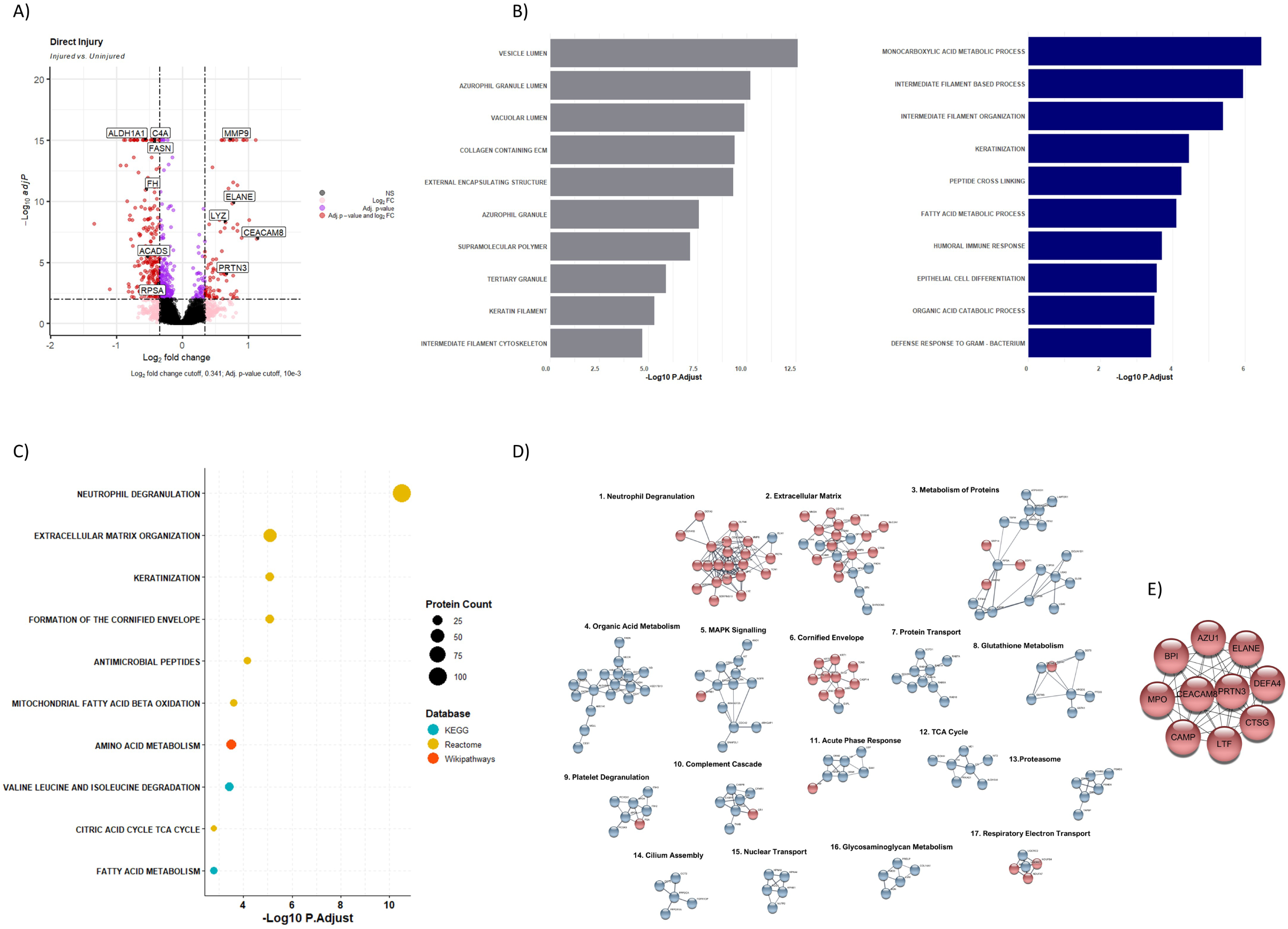
Direct (pulmonary) injury profile: A) Volcano plot of direct injury comparison between injured group vs uninjured group. X-axis represents the log2 foldchange while the y-axis the -log10 q value. Circles in red were significant based on cut-off while pink had changed in log2 foldchange but not g-value while blue had a significant g-value but no change in the log2foldchange. B) Bar chart of direct injury comparison functional annotation using GO, grouped by category grey representing cellular component and blue biological process. X-axis -log10 adjusted p value for the pathways tested. C) Dot plot of direct injury comparison pathway analysis results, color of dot represents source database, and size represents number of proteins within that pathway. X-axis -log10 adjusted p value for the pathways tested. D) Functional clusters within direct injury comparison PPl network with 25 proteins. E) Direct injury comparison PPI network of the hub genes identified.

Protein–protein interaction analysis using STRING in Cytoscape identified interacting modules annotated as neutrophil degranulation, extracellular matrix organisation, metabolic reprogramming and MAPK signalling, with neutrophil granule-associated proteins among the top network hubs (Fig. 4D–E). Cell-type enrichment prioritised epithelial cells, monocytes and neutrophils (Supplementary Fig. 1A).

### Indirect lung injury programs

#### Indirect injury features complement/coagulation cascade perturbation with stress and translation-linked network architecture

In indirect injury differential expression analysis using MS_EmpiRe revealed 394 proteins differed between injured and matched uninjured controls (Fig. 5A), comprising 252 differentially expressed proteins (87 increased; 165 decreased) and 142 differentially detected proteins (Logfold 0.347 adj p <0.01). Functional enrichment was dominated by complement and coagulation cascades, acute phase/wound healing programs and extracellular matrix organisation (Fig. 5B), with canonical pathway analysis highlighting complement/coagulation cascade perturbation alongside ECM organisation programs (Fig. 5C). Protein–protein interaction analysis produced a network with 394 nodes and 661 edges and identified major clusters including coagulation, complement activation, extracellular matrix, translation-related modules and oxidative phosphorylation (Fig. 5D). To identify the most highly connected nodes within the protein-protein interaction network we used cytoHubba in Cytoscape to identify the top ten highest ranking nodes. These hub proteins, were predominantly ribosomal proteins, consistent with translation-linked regulation within the indirect injury network (Fig. 5E). Cell-type enrichment prioritised barrier-associated populations including epithelial, endothelial, fibroblast and stromal cells (Supplementary Fig. 1B).

**Figure 5:**
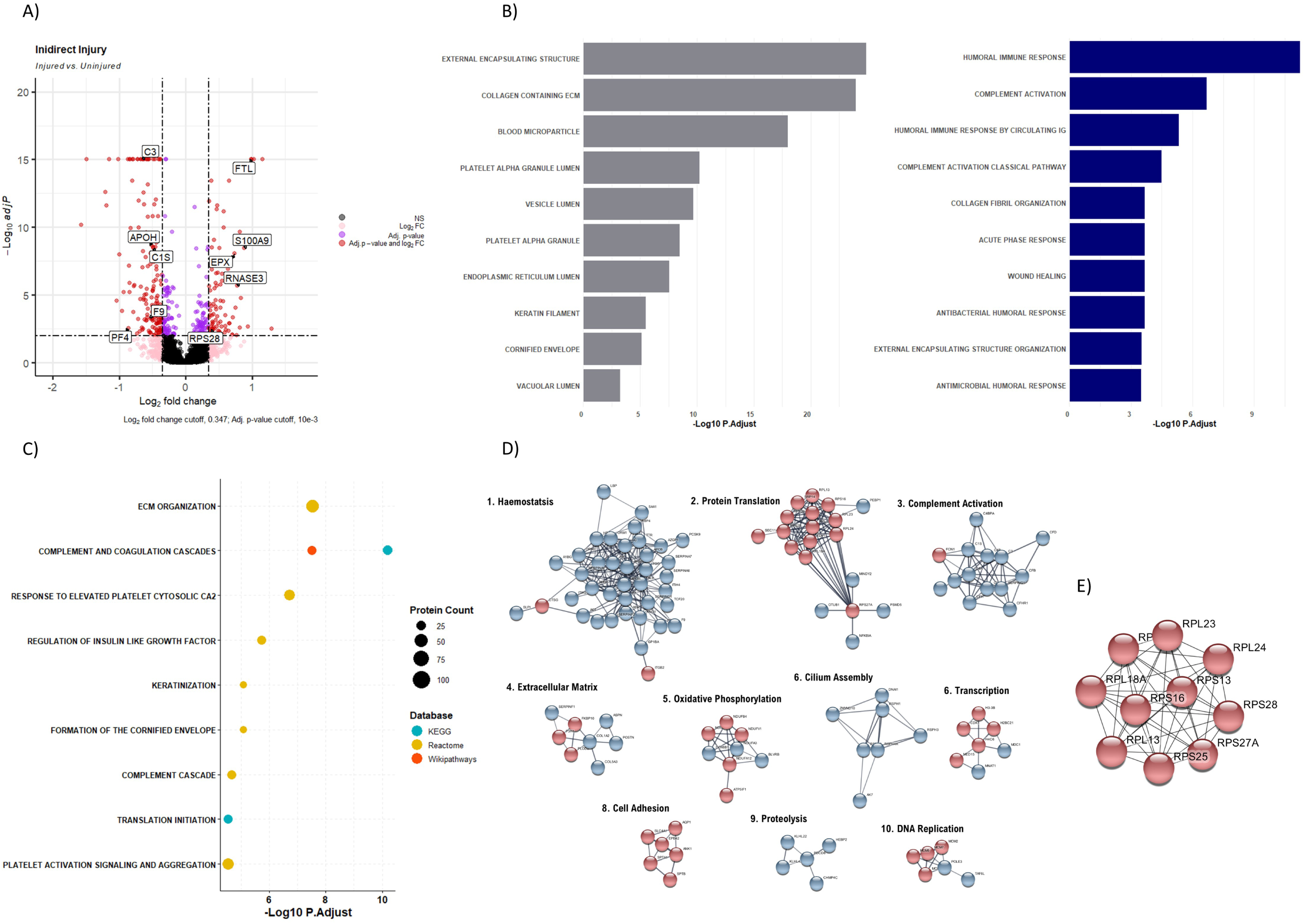
Indirect (extra-pulmonary) injury profile: A) Volcano plot of indirect injury comparison between injured group vs uninjured group. X-axis represents the log2 foldchange while the y-axis the -log10 q value. Circles in red were significant based on cut-off while pink had changed in log2 foldchange but not g-value while blue had a significant q-value but no change in the log2foldchange. B) Bar chart of indirect injury comparison functional annotation using GO, grouped by category grey representing cellular component and blue biological process. X-axis -log10 adjusted pv alue for the pathways evaluated. C) Dot plot of indirect injury comparison pathway analysis results, color of dot represents source database and size represents number of proteins within that pathway. X-axis -log10 adjusted p value for the pathways. D) Functional clusters within indirect injury comparison PPl network with 25 proteins E} Indirect injury comparison PP network of the hub genes identified.

### EVLP platform-associated proteomic effects

To quantify platform-associated biology, baseline tissue was compared with an uninjured segment at end-EVLP (Supplementary Fig. 2A). Differential testing identified 413 proteins altered by EVLP conditions alone (116 differentially expressed; 293 differentially detected). Enrichment analyses highlighted extracellular matrix organisation, hemostasis/complement programs, acute phase responses and lipid/cholesterol metabolism modules (Supplementary Fig. 2B–E). Cell-type enrichment for the EVLP-effect signature is shown in Supplementary Fig. 1C. These EVLP-associated shifts support the use of within-lung controls and explicit EVLP-effect contrasts when interpreting injury signatures and designing EVLP interventional studies.

### Therapeutic discovery

Therapeutic nomination and cross-platform prioritisation were anchored on proteomics-derived end-EVLP injured vs uninjured contrasts (direct: injured vs uninjured segment; indirect: injured vs matched uninjured segment). We nominated repurposing candidates by querying proteomics-derived injury signatures against the CLUE L1000 perturbational compendium and selecting perturbagens with significant negative connectivity (FDR < 0.05). We then prioritised candidates by cross-platform concordance (Table 1; Supplementary Tables 2–3).

**Table 1.**
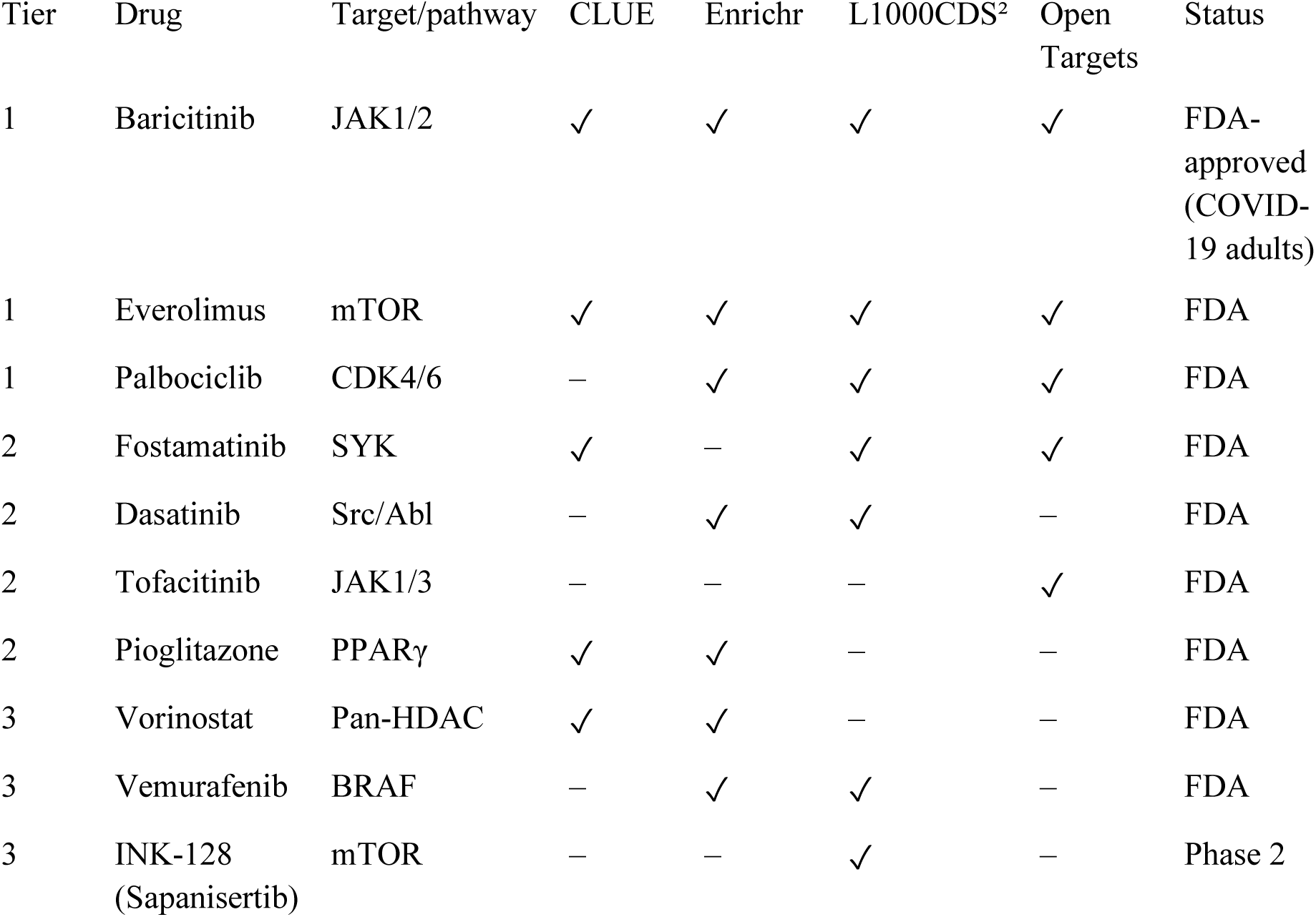
Cross-platform tiered prioritisation of candidate therapeutics for EVLP validation Tiered prioritisation derived from concordant evidence across CLUE L1000, Enrichr, L1000CDS? and Open Targets. Tick marks indicate supportive evidence for the nominated class/agent in the cross platform validation.

| Tier | Drug | Target/pathway | CLUE | Enrichr | L1000CDS² | Open Targets | Status |
|---|---|---|---|---|---|---|---|
| 1 | Baricitinib | JAK1/2 | ✓ | ✓ | ✓ | ✓ | FDA-approved (COVID-19 adults) |
| 1 | Everolimus | mTOR | ✓ | ✓ | ✓ | ✓ | FDA |
| 1 | Palbociclib | CDK4/6 | – | ✓ | ✓ | ✓ | FDA |
| 2 | Fostamatinib | SYK | ✓ | – | ✓ | ✓ | FDA |
| 2 | Dasatinib | Src/Abl | – | ✓ | ✓ | – | FDA |
| 2 | Tofacitinib | JAK1/3 | – | – | – | ✓ | FDA |
| 2 | Pioglitazone | PPARγ | ✓ | ✓ | – | – | FDA |
| 3 | Vorinostat | Pan-HDAC | ✓ | ✓ | – | – | FDA |
| 3 | Vemurafenib | BRAF | – | ✓ | ✓ | – | FDA |
| 3 | INK-128 (Sapanisertib) | mTOR | – | – | ✓ | – | Phase 2 |

### Cross-platform validation of therapeutic candidates

To reduce dependence on any single perturbational resource, we cross-validated CLUE-nominated therapeutic axes using Enrichr, L1000CDS² and the Open Targets Platform. Across platforms, recurrent signals centred on PI3K/AKT/mTOR, CDK, SYK and JAK/STAT pathways, with additional support for HDAC inhibition and PPARγ/metabolic restoration. These convergent axes defined a tiered shortlist for EVLP intervention testing (Table 1; Supplementary Tables 2–3).

## Discussion

This study establishes a paired human EVLP injury platform that resolves early direct versus indirect lung injury biology and explicitly links these mechanisms to therapeutic hypotheses using signature-reversal analysis. Three findings are particularly relevant to therapeutic discovery.

First, direct injury rapidly engages neutrophil granule biology and tissue remodelling programs, consistent with canonical roles of neutrophil proteases in barrier disruption and extracellular matrix degradation (12,40–42). Notably, this was seen under acellular perfusion consistent with activation of a pre-existing lung neutrophil pool. In this context, the timing of intervention is likely critical: inhibitors targeting neutrophil effector proteins have shown mixed clinical efficacy, potentially because treatment is administered after early neutrophil-driven injury has already occurred (43). The direct injury proteome also highlighted metabolic rewiring in early injury, consistent with emerging work linking lipid and fatty acid metabolism to acute lung injury and immunometabolism (44–45). Chemokine-dependent neutrophil recruitment and synergy may further amplify direct injury programs (46).

Second, indirect injury preferentially perturbs complement and coagulation cascades alongside stress-response and translation-associated programs, consistent with immunothrombosis and systemic inflammatory-coagulant crosstalk described in ARDS (14,47–49). These observations align with complement and anticoagulant/immunomodulatory strategies proposed for ARDS and sepsis-associated lung injury (50–53). Notably, ribosomal proteins formed hub nodes within the indirect injury network, consistent with reported extraribosomal roles for select ribosomal proteins in regulating inflammatory resolution and stress responses (including GAIT-complex function and p53 pathway modulation) (54–57).

Third, signature reversal analysis and cross-platform validation converged on druggable pathway axes including JAK/STAT, PI3K/AKT/mTOR, SYK and CDK programs, with supportive evidence for HDAC inhibition. Genetic/disease association evidence supports prioritisation of JAK/STAT-, SYK- and PI3K/mTOR-linked targets in respiratory disease contexts. As a translational precedent for host-directed signalling modulation in viral pneumonia-associated ARDS, JAK1/2 inhibition with baricitinib improved outcomes in hospitalised COVID-19 in randomised trials (58–59). These convergent signals support pathway-level rather than single-compound prioritisation and provide a rational short-list for EVLP intervention experiments that test both physiologic rescue and reversal of injury associated proteomic signatures.

EVLP itself induced extracellular matrix, hemostasis and lipid metabolism shifts, emphasising the need to control for platform-associated biology. These EVLP-associated changes are consistent with ischemia-reperfusion-related inflammatory states described during ex vivo perfusion (60).

Several limitations should be considered. EVLP does not recapitulate whole-body immune and neurohumoral interactions, which may particularly affect indirect injury. The 4 h timepoint captures early injury but cannot resolve later-phase transitions. Finally, CLUE L1000 predictions derive from cancer cell line perturbations (here A549 and HCC515) and should be viewed as hypothesis-generating rather than proof of efficacy. Collectively, these limitations motivate direct EVLP intervention experiments as the next step.

Whole-organ EVLP offers a unique translational bridge between reductionist human systems (cells, organoids and precision-cut lung slices) and the complexity of clinical ARDS. Whereas organoids and cell culture models provide tractable mechanistic perturbation, they typically lack a perfused microvasculature, intact alveolar–capillary geometry and resident immune microenvironments, limiting interrogation of endothelial–epithelial cross-talk and compartmental injury (61). Precision-cut lung slices preserve native extracellular matrix and multicellular architecture and are powerful for higher-throughput human tissue experimentation, but cannot reproduce whole-organ ventilation–perfusion coupling or deliver matched airway-versus-vascular insults within the same lung (62). By maintaining an intact bronchial tree, interstitium and microvascular bed under controlled ventilation and perfusion, our EVLP platform enables direct (endobronchial) versus indirect (intravascular) injury to be applied within paired donor lungs, allowing compartment-specific molecular programmes to be resolved with reduced inter-donor noise.

A notable feature of this whole-lung preparation is the persistence of resident and marginated leukocyte pools within the pulmonary microvasculature and interstitium at the time of experimentation. The lung microvasculature is increasingly recognised as an immune niche enriched with marginated, primed and aged neutrophils poised for rapid responses to blood-borne stimuli (63). Consistent with this biology, marginated neutrophils have been shown to remain functionally relevant in ex vivo perfused human lungs (64). These observations provide an explanation for the prominent neutrophil degranulation signature and MPO+ staining observed here despite the use of an acellular perfusate, and highlight an under-appreciated strength of EVLP for modelling the earliest, tissue-resident phase of neutrophil-mediated injury without confounding by ongoing systemic recruitment. Complementary in situ human lung imaging approaches— including bedside optical endomicroscopy of activated neutrophils (65) and ex vivo lung ventilation– based immuno-imaging of resident lymphocyte populations (66)—support the feasibility of spatially resolving resident immune programmes within intact human lungs. Conversely, the absence of circulating blood cells remains a limitation for modelling later recruitment and immunothrombotic amplification, motivating future experiments that incorporate cellular perfusates or staged add-back approaches.

In summary, this study couples a paired human EVLP injury platform with proteomic profiling and signature-based therapeutic discovery. The resulting injury-stratified hypotheses provide a rational basis for prioritised validation studies within EVLP and subsequently in translational models, moving toward biologically matched interventions for ARDS (67).

## Data availability

The mass spectrometry proteomics data have been deposited to the ProteomeXchange Consortium via the PRIDE partner repository under accession code PXD087977 and Project DOI: 10.6019/PXD087977. The dataset is private during peer review and is available to editors and reviewers using the credentials provided in the confidential submission material.

## Methods

### Ethics and human lung procurement

Human donor lungs declined for transplantation were procured from the International Institute for the Advancement of Medicine (Edison, NJ) following approval from the appropriate regional ethics committee and with informed consent from donors or their next of kin.

### Ex vivo lung perfusion (EVLP) platform

Single-lung EVLP experiments were conducted. Where feasible, paired direct (endobronchial) and indirect (perfusate) injury models were performed using lungs from the same donor. An acellular perfusate was used, consisting of 2 L Dulbecco’s Modified Eagle’s Medium (DMEM; high glucose), 10,000 IU heparin, 7% bovine serum albumin and 10 g dextran-40. Perfusion was started at 0.2 L/min and maintained between 0.2–0.5 L/min throughout experiments to ensure pulmonary artery pressure remained <10 mmHg. Mechanical ventilation was initiated once organ temperature reached 32 °C (10 breaths/min, tidal volume 2.5–3 mL/kg, positive end-expiratory pressure 5 cmH2O).

### Injury induction

Direct injury: lipopolysaccharide (LPS; Escherichia coli O111:B4) 6 mg in 10 mL normal saline was delivered endobronchially to a subsegment of the right lower lobe under bronchoscopic guidance.

Indirect injury: LPS 5 μg/kg ideal body weight (dose range 300–800 μg) diluted in 10 mL sterile normal saline was infused into the pulmonary artery limb of the circuit over 5 minutes to model endotoxemia.

### Tissue sampling and experimental design

Tissue samples were collected at baseline (start of mechanical ventilation) and at the end of the experiment (4 h post-LPS). In the endobronchial LPS model, baseline and injured samples were obtained from the right lower lobe, and uninjured control samples from the right upper lobe. In the perfusate LPS model, baseline and injured samples were obtained from the left lower lobe. Tissue (6–9 cm3) was collected using a handheld biopsy stapler (Endo GIA; Covidien), and either snap-frozen for proteomics or fixed for histology. Perfusate samples were collected at circuit set-up, baseline and hourly after LPS administration.

### Histology and lung injury scoring

Lung tissues were fixed in 10% neutral buffered formalin, processed and embedded in paraffin. Sections (5 μm) were stained with hematoxylin and eosin (H&E) and digitally scanned. Lung injury scoring was performed using established criteria as previously reported (ATS experimental acute lung injury workshop report).

### Tissue microarray (TMA) and neutrophil quantification

All slides were reviewed by an expert pathologist and three representative areas were selected per sample. A 1 mm punch was taken from each donor block and embedded to create tissue microarray blocks representative of the cohort. Sections were stained with anti-myeloperoxidase (MPO) primary antibody and Vector® Red AP substrate. MPO-positive cells were quantified as an index of neutrophil infiltration.

### Perfusate cytokines and adhesion molecules

Perfusate cytokines and soluble adhesion molecules were measured at circuit set-up, baseline and hourly following LPS administration. Tumour necrosis factor alpha (TNFα), interleukin-1 beta (IL-1β), interleukin-6 (IL-6), interleukin-8 (IL-8), soluble ICAM-1 (sICAM-1) and soluble VCAM-1 (sVCAM-1) were quantified using the Simple Plex™ Ella microfluidic automated ELISA platform (ProteinSimple, CA, USA) according to the manufacturer’s instructions.

### Proteomics sample processing

Snap-frozen tissue was extracted in denaturing buffer, homogenised and sonicated. Tryptic digestion was performed using the S-Trap method. Peptides were analysed by liquid chromatography–mass spectrometry (LC–MS/MS) on a timsTOF FleX mass spectrometer at the Proteomics and Metabolomics Facility at the Roslin Institute, University of Edinburgh.. Raw data were processed using Spectronaut and DIA-NN.

### Proteomic data processing and differential analysis

Spectra acquired on a timsTOF FleX were processed in Spectronaut and analysed with MS-DAP (v1.0.6) under R 4.2.1. Precursors were mapped to the human reference proteome (UP000005640_9606). Peptides were retained when detected and quantified in at least three samples and ≥75% of samples within each group, with ≥2 peptides per protein, using MS-DAP’s global (“by dataset”) filter; intensities were normalised by vsn followed by mode-between-protein normalisation and rolled up to protein abundances using MaxLFQ. Differential expression was assessed with MS-EmpiRe across three pre-specified contrasts: the EVLP effect (endobronchial-LPS baseline versus uninjured segment), direct injury (uninjured versus endobronchial-LPS segment) and indirect injury (uninjured segment versus perfusate-LPS lung). Proteins were considered significant at FDR-adjusted q < 0.01 with an absolute log2 fold-change exceeding a per-contrast threshold estimated by MS-DAP’s bootstrap procedure (0.307, 0.341 and 0.347, respectively). Proteins lacking sufficient peptide-level data for differential expression were additionally evaluated by differential detection from observed peptide counts (≥2 peptides in ≥3 samples and ≥50% of replicates per group); proteins with an absolute differential-detection z-score ≥ 5 were retained. For each injury contrast, the differentially expressed and differentially detected proteins were combined by direction of change to define the injury signature used for downstream enrichment and connectivity analyses.

### Dimensionality reduction and clustering

Protein-level MaxLFQ abundances were scaled to unit variance, the lowest-variance 10% of proteins removed, and principal component analysis performed with PCAtools (v2.8.0), displayed with 75% t-distributed confidence ellipses by group. For visualisation of injury signatures, significant-protein abundances were row z-scored and clustered by hierarchical clustering (Euclidean distance, Ward.D linkage) using pheatmap, partitioned into four modules.

### Functional enrichment

Over-representation analysis was performed per contrast on the injury-signature gene lists in clusterProfiler (v4.4.4) using enricher against MSigDB v2023.2.Hs gene sets (C2:CP Reactome, C2:CP canonical and C2:CP KEGG-legacy) and enrichGO (Biological Process and Cellular Component ontologies; org.Hs.eg.db v3.15.0), with the detected proteome as the background universe. Terms were considered enriched at Benjamini–Hochberg-adjusted p < 0.05 with more than five member genes.

### Protein–protein interaction networks

Interaction networks were constructed from the injury-signature proteins in STRING v12.0 (stringApp v2.0.3) within Cytoscape v3.10.1 at a minimum interaction confidence of 0.4. Functional modules were identified by Markov Cluster (MCL) clustering (clusterMaker2; granularity/inflation 2.5) and annotated against KEGG, Reactome and WikiPathways. Hub proteins were defined as the ten highest-ranked nodes by Maximal Clique Centrality (MCC) using cytoHubba (v0.1).

### Cell-type enrichment

The combined differential-expression and differential-detection gene lists for each contrast were submitted to WebCSEA (Dai et al., 2022; https://bioinfo.uth.edu/webcsea/, accessed 23 February 2024). General cell types were ranked by the WebCSEA combined p-value; enrichment was considered significant at a Bonferroni-corrected threshold of p < 3.69 × 10^⁻□^ (correcting across the 1,355 tissue-cell types in the reference panel) and at a nominal threshold of p < 1 × 10^-3^. The 20 highest-ranked general cell types per contrast are reported.

### Cross-platform drug repurposing validation

Therapeutic repurposing candidates were cross-validated using four computational resources applied to the same EVLP-derived signatures: (1) CLUE L1000 (Broad Connectivity Map) using connectivity scoring across LINCS L1000 perturbational profiles; (2) Enrichr (Ma’ayan Laboratory) using Fisher’s exact test enrichment across multiple drug perturbation libraries; (3) L1000CDS² using the characteristic direction algorithm to identify perturbagens whose signatures oppose the query; and (4) Open Targets Platform to evaluate genetic/genomic evidence linking nominated targets to respiratory and inflammatory diseases.

Input signatures were defined from the EVLP proteomic analyses and separated by injury model and direction of change: Direct UP (99), Direct DOWN (358), Indirect UP (91), and Indirect DOWN (303). Reversal logic was defined as disease UP + drug DOWN and disease DOWN + drug UP. Where gene lists exceeded the CLUE 150-gene input limit, signatures were truncated to the 150 most differentially expressed proteins ranked by absolute fold change. For downregulated signatures, the 10 most upregulated proteins from the corresponding contrast were included as the required upregulated anchor. Candidate classes were prioritised based on recurrence across platforms and, where applicable, availability of clinically approved compounds. In addition to signalling-node inhibitors, epigenetic modulators (including HDAC inhibitors) were captured by this approach via both lung-context connectivity in CLUE and broad perturbation-library enrichment in Enrichr.

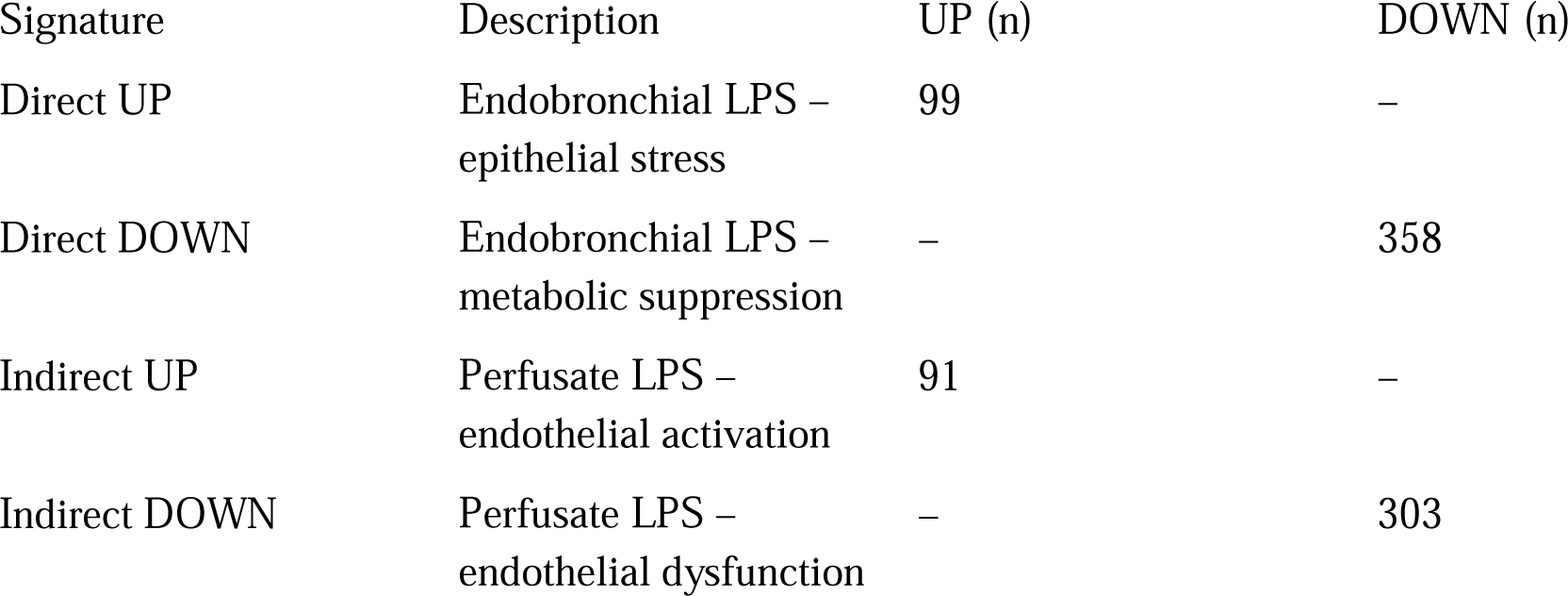

### Therapeutic connectivity analysis (CLUE/L1000)

Proteomics-derived injury signatures were defined from within-lung, end-EVLP contrasts (direct injured vs uninjured segment; indirect injured vs matched uninjured segment) and queried against the CLUE L1000 perturbational database. Compounds with statistically significant negative connectivity (FDR < 0.05) were interpreted as predicted reversers of injury signatures. CLUE queries were run against all cell lines; results were additionally filtered to A549 and HCC515 for lung-specific prioritisation

---

**Supplementary Figure 1:**
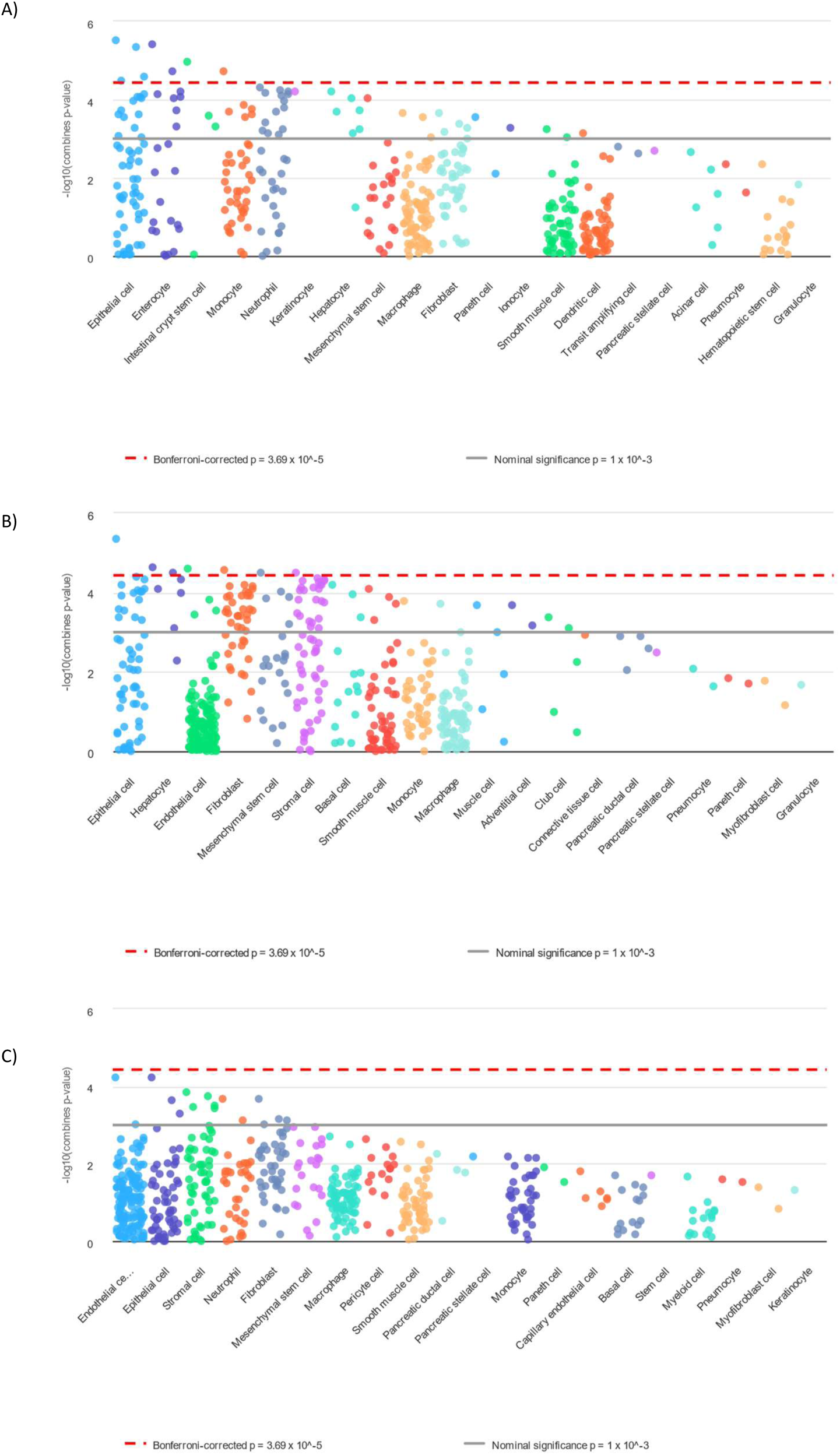
Cell-type enrichment using WebSCEA: Manhattan plot of the top 20 cell types over enriched for differentially identified proteins in: A) Direct injury. B) Indirect injury. C) EVLP comparison

**Supplementary figure 2:**
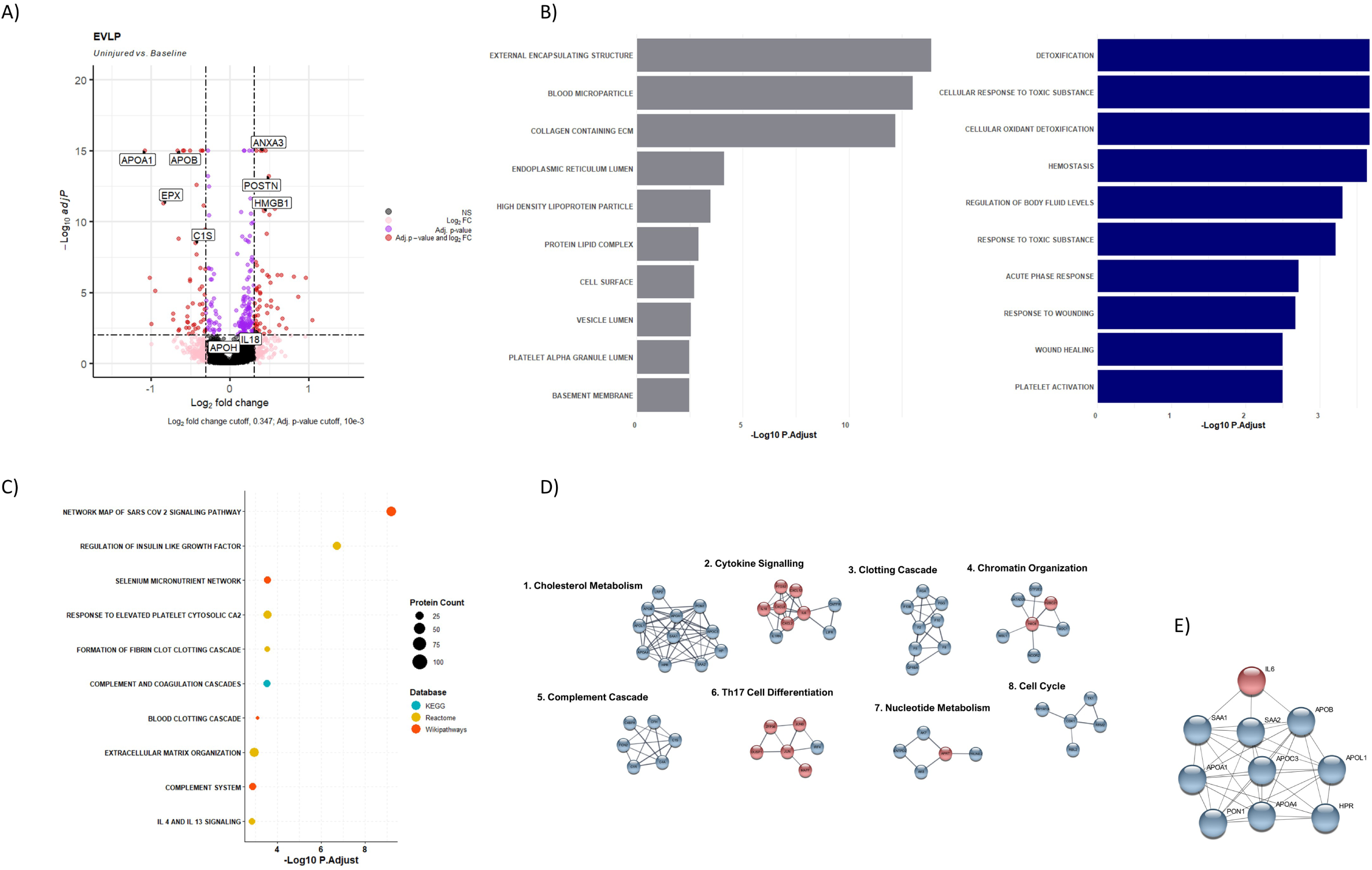
Effect of EVLP: A) Volcano plot of EVLP comparison between uninjured group vs baseline group. X-axis represents the log2 foldchange while the y-axis the -log10 q value. Circles in red were significant based on cut-off while pink had changed in log2 foldchange but not g-value while blue had a significant g-value but no change in the log2foldchange. B) Bar chart of EVLP proteins functional annotation using GO, grouped by category grey representing cellular component and blue biological process. X-axis -log10 adjusted p value for the pathways tested. C) Dot plot of EVLP proteins pathway analysis results, color of dot represents source database and size represents number of proteins within that pathway. X-axis -log10 adjusted p value for the pathways tested. D) Functional clusters within EVLP PPI network with 25 proteins. E) EVLP PPl network of the hub genes identified.

**Supplementary Table 1.**
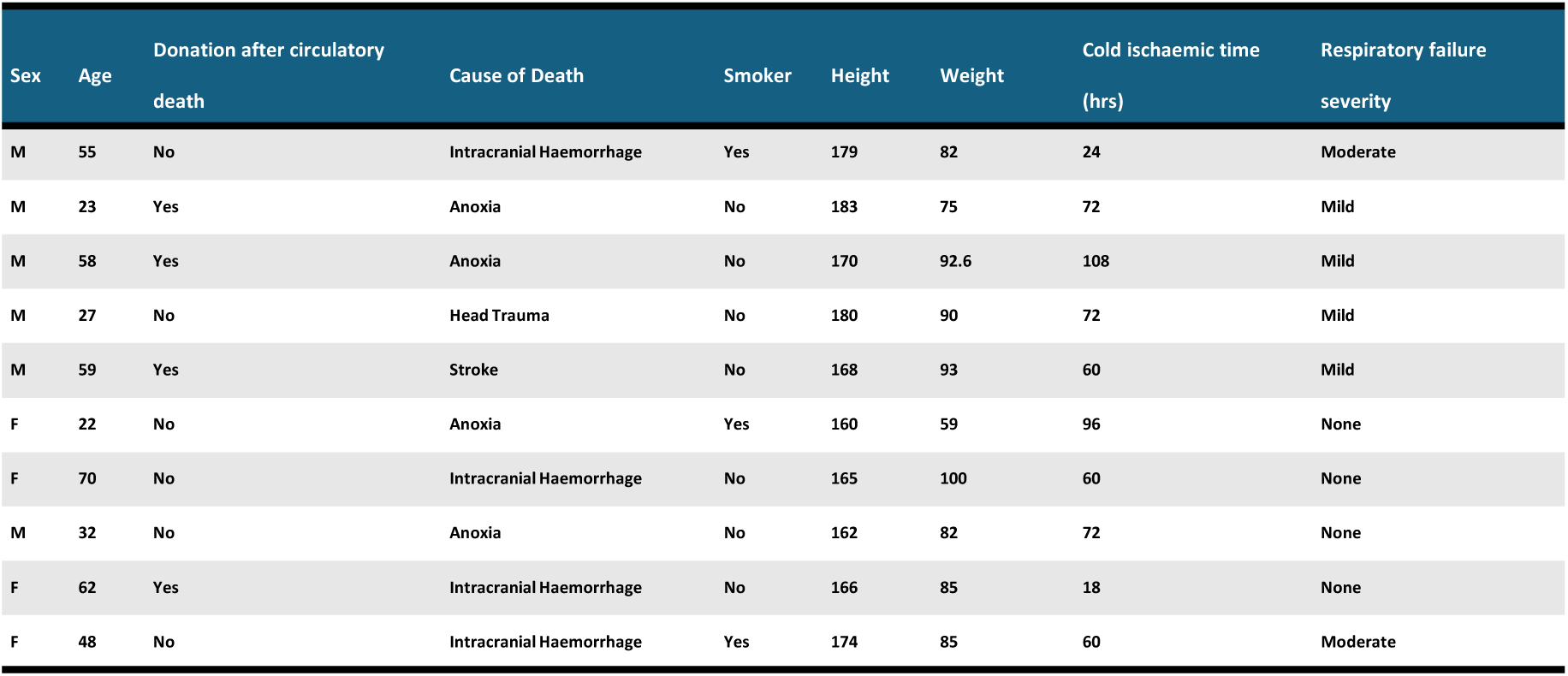
Clinical details of donors.

**Supplementary Table 2.**
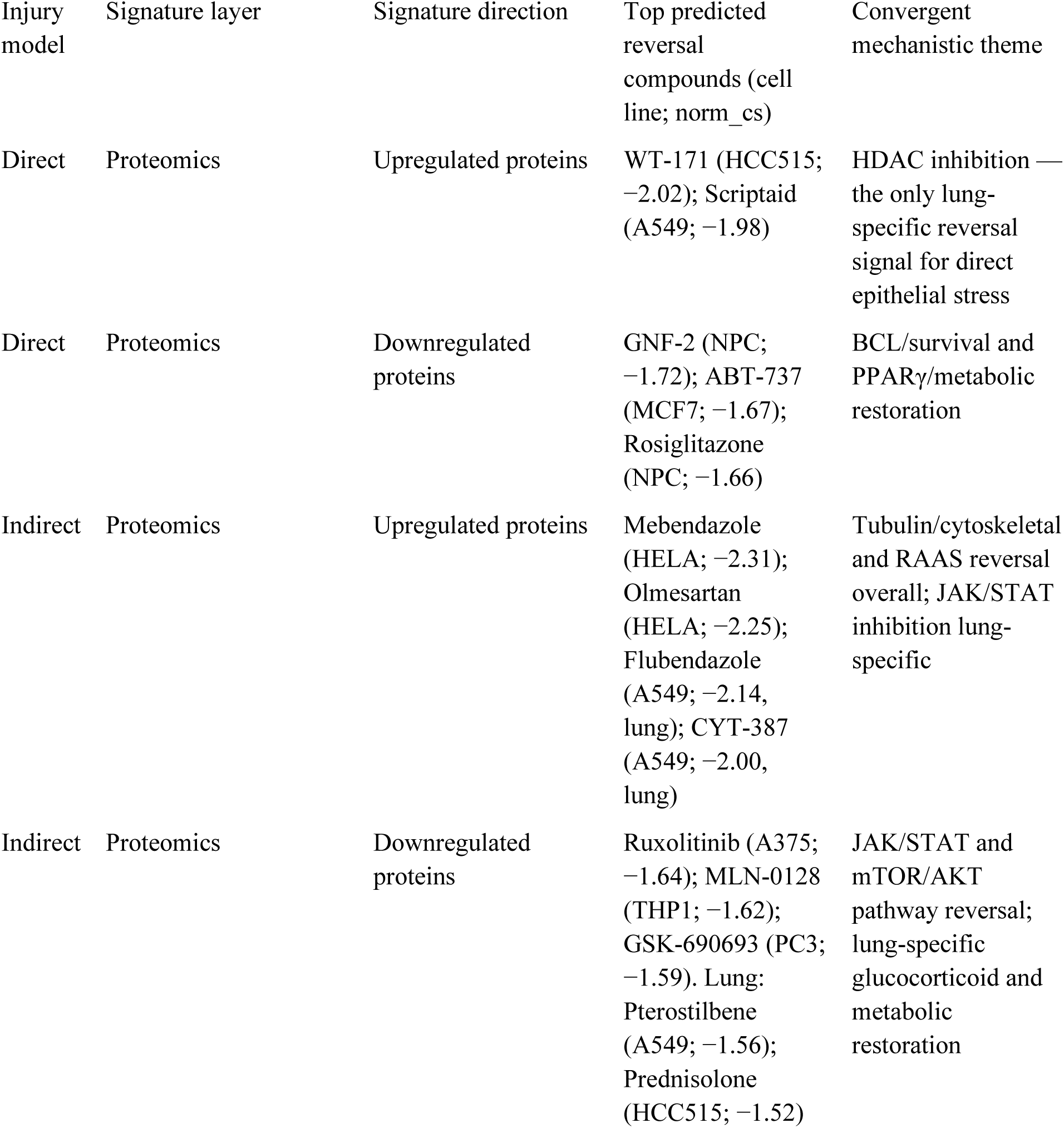
Therapeutic signature reversal highlights injury-specific candidate drug Classes.

**Supplementary Table 3.**
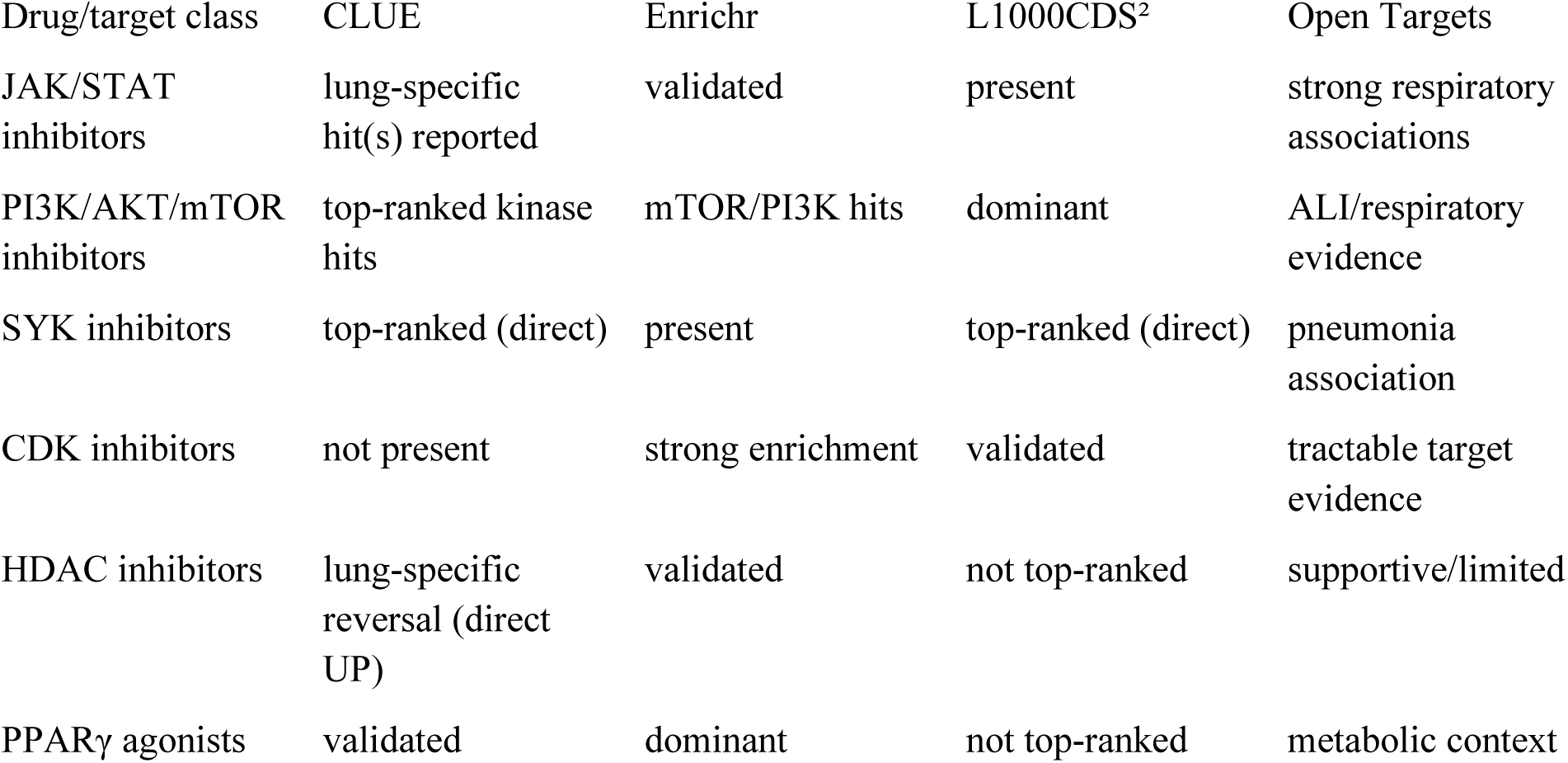
Cross-platform class concordance summary.

## Notes

### Competing Interest Statement

The authors have declared no competing interest.

### Summary of Updates

Adding PRIDE data repository accession codes for proteomics data to the data availability statement.

## References

1. Bellani, G. et al. Epidemiology, patterns of care, and mortality for patients with acute respiratory distress syndrome in intensive care units in 50 countries. JAMA 315, 788–800 (2016). doi:10.1001/jama.2016.0291

2. Herridge, M. S. et al. Functional disability 5 years after acute respiratory distress syndrome. N. Engl. J. Med. 364, 1293–1304 (2011). doi:10.1056/NEJMoa1011802

3. Wilson, J. G. & Calfee, C. S. ARDS subphenotypes: understanding a heterogeneous syndrome. Crit. Care 24, 102 (2020). doi:10.1186/s13054-020-2778-x

4. Matthay, M. A., McAuley, D. F. & Ware, L. B. Clinical trials in acute respiratory distress syndrome: challenges and opportunities. Lancet Respir. Med. 5, 524–534 (2017). doi:10.1016/S2213-2600(17)30188-1

5. Calfee, C. S. et al. Distinct molecular phenotypes of direct vs indirect ARDS in single-center and multicenter studies. Chest 147, 1539–1548 (2015). doi:10.1378/chest.14-2454

6. Gattinoni, L. et al. Acute respiratory distress syndrome caused by pulmonary and extrapulmonary disease. Different syndromes? Am. J. Respir. Crit. Care Med. 158, 3–11 (1998). doi:10.1164/ajrccm.158.1.9708031

7. Tugrul, S. et al. Effects of sustained inflation and postinflation positive end-expiratory pressure in acute respiratory distress syndrome: focusing on pulmonary and extrapulmonary forms. Crit. Care Med. 31, 738–744 (2003). doi:10.1097/01.CCM.0000053554.76355.72

8. Maron-Gutierrez, T. et al. Effects of mesenchymal stem cell therapy on the time course of pulmonary remodeling depend on the etiology of lung injury in mice. Crit. Care Med. 41, e319–e333 (2013). doi:10.1097/CCM.0b013e31828a663e

9. Uhlig, C. et al. The effects of salbutamol on epithelial ion channels depend on the etiology of acute respiratory distress syndrome but not the route of administration. Respir. Res. 15, 56 (2014). doi:10.1186/1465-9921-15-56

10. Leite-Junior, J. H. et al. Methylprednisolone improves lung mechanics and reduces the inflammatory response in pulmonary but not in extrapulmonary mild acute lung injury in mice. Crit. Care Med. 36, 2621–2628 (2008). doi:10.1097/CCM.0b013e3181847b43

11. Matthay, M. A. et al. Acute respiratory distress syndrome. Nat. Rev. Dis. Primers 5, 18 (2019). doi:10.1038/s41572-019-0069-0

12. Potey, P. M. D., Rossi, A. G., Lucas, C. D. & Dorward, D. A. Neutrophils in the initiation and resolution of acute pulmonary inflammation: understanding biological function and therapeutic potential. J. Pathol. 247, 672–685 (2019). doi:10.1002/path.5221

13. Schmidt, E. P. et al. The circulating glycosaminoglycan signature of respiratory failure in critically ill adults. J. Biol. Chem. 289, 8194–8202 (2014). doi:10.1074/jbc.M113.539452

14. Frantzeskaki, F., Armaganidis, A. & Orfanos, S. E. Immunothrombosis in acute respiratory distress syndrome: cross talks between inflammation and coagulation. Respiration 93, 212–225 (2017). doi:10.1159/000453002

15. Mir, T. et al. Outcome and post-surgical lung biopsy change in management of ARDS: a proportional prevalence meta-analysis. Adv. Respir. Med. 90, 267–278 (2022). doi:10.3390/arm90040036

16. Bowler, R. P. et al. Proteomic analysis of pulmonary edema fluid and plasma in patients with acute lung injury. Am. J. Physiol. Lung Cell Mol. Physiol. 286, L1095–L1104 (2004). doi:10.1152/ajplung.00304.2003

17. de Torre, C. et al. Proteomic analysis of inflammatory biomarkers in bronchoalveolar lavage. Proteomics 6, 3949–3957 (2006). doi:10.1002/pmic.200500693

18. Bhargava, M. et al. Proteomic profiles in acute respiratory distress syndrome differentiates survivors from non-survivors. PLoS One 9, e109713 (2014). doi:10.1371/journal.pone.0109713

19. Chen, X. et al. Quantitative proteomic analysis by iTRAQ for identification of candidate biomarkers in plasma from acute respiratory distress syndrome patients. Biochem. Biophys. Res. Commun. 441, 1–6 (2013). doi:10.1016/j.bbrc.2013.09.027

20. Chang, D. W. et al. Proteomic and computational analysis of bronchoalveolar proteins during the course of the acute respiratory distress syndrome. Am. J. Respir. Crit. Care Med. 178, 701–709 (2008). doi:10.1164/rccm.200712-1895OC

21. Morrell, E. D. et al. Peripheral and alveolar cell transcriptional programs are distinct in acute respiratory distress syndrome. Am. J. Respir. Crit. Care Med. 197, 528–532 (2018). doi:10.1164/rccm.201703-0614LE

22. Kang, Z. Y. et al. Heterogeneity of immune cells and their communications unveiled by transcriptome profiling in acute inflammatory lung injury. Front. Immunol. 15, 1382449 (2024). doi:10.3389/fimmu.2024.1382449

23. Frohlich, E. Animals in respiratory research. Int. J. Mol. Sci. 25, 2903 (2024). doi:10.3390/ijms25052903

24. Abdalla, A., Dhaliwal, K. & Shankar-Hari, M. Ex vivo lung perfusion models to explore the pathobiology of ARDS. In Annual Update in Intensive Care and Emergency Medicine 2023 (ed. Vincent, J.-L.) 111–119 (Springer, 2023). doi:10.1007/978-3-031-23005-9_9

25. Lee, J. W. et al. Allogeneic human mesenchymal stem cells for treatment of E. coli endotoxin-induced acute lung injury in the ex vivo perfused human lung. Proc. Natl Acad. Sci. USA 106, 16357–16362 (2009). doi:10.1073/pnas.0907996106

26. Weathington, N. M. et al. Ex vivo lung perfusion as a human platform for preclinical small molecule testing. JCI Insight 3, e95515 (2018). doi:10.1172/jci.insight.95515

27. Roffia, V. et al. Proteome investigation of rat lungs subjected to ex vivo perfusion (EVLP). Molecules 23, 3061 (2018). doi:10.3390/molecules23123061

28. Niroomand, A. et al. Proteomic changes to immune and inflammatory processes underlie lung preservation using ex vivo cytokine adsorption. Front. Cardiovasc. Med. 10, 1274444 (2023). doi:10.3389/fcvm.2023.1274444

29. Beutler, B. & Rietschel, E. T. Innate immune sensing and its roots: the story of endotoxin. Nat. Rev. Immunol. 3, 169–176 (2003). doi:10.1038/nri1004

30. Imai, Y. et al. Identification of oxidative stress and Toll-like receptor 4 signaling as a key pathway of acute lung injury. Cell 133, 235–249 (2008). doi:10.1016/j.cell.2008.02.043

31. Kawai, T. & Akira, S. TLR signaling. Cell Death Differ. 13, 816–825 (2006). doi:10.1038/sj.cdd.4401850

32. Matute-Bello, G. et al. An official American Thoracic Society workshop report: features and measurements of experimental acute lung injury in animals. Am. J. Respir. Cell Mol. Biol. 44, 725–738 (2011). doi:10.1165/rcmb.2009-0210ST

33. Koopmans, F., et al. MS-DAP platform for downstream data analysis of label-free proteomics uncovers optimal workflows in benchmark data sets and increased sensitivity in analysis of Alzheimer’s biomarker data. J. Proteome Res. 22, 374–386 (2023). doi:10.1021/acs.jproteome.2c00513

34. Wu, T. et al. clusterProfiler 4.0: a universal enrichment tool for interpreting omics data. Innovation 2, 100141 (2021). doi:10.1016/j.xinn.2021.100141

35. Kanehisa, M. & Goto, S. KEGG: Kyoto encyclopedia of genes and genomes. Nucleic Acids Res. 28, 27–30 (2000). doi:10.1093/nar/28.1.27

36. Martens, M., et al. WikiPathways: connecting communities. Nucleic Acids Res. 49, D613–D621 (2021). doi:10.1093/nar/gkaa1024

37. Milacic, M. et al. The Reactome pathway knowledgebase 2024. Nucleic Acids Res. 52, D672–D678 (2024). doi:10.1093/nar/gkad1025

38. Chin, C.-H. et al. cytoHubba: identifying hub objects and sub-networks from complex interactome. BMC Syst. Biol. 8 (Suppl 4), S11 (2014). doi:10.1186/1752-0509-8-S4-S11

39. Dai, Y. et al. WebCSEA: web-based cell-type-specific enrichment analysis of genes. Nucleic Acids Res. 50, W782–W790 (2022). doi:10.1093/nar/gkac392

40. Korkmaz, B. et al. Neutrophil elastase, proteinase 3, and cathepsin G as therapeutic targets in human diseases. Pharmacol. Rev. 62, 726–759 (2010). doi:10.1124/pr.110.002733

41. Boxio, R. et al. Neutrophil elastase cleaves epithelial cadherin in acutely injured lung epithelium. Respir. Res. 17, 129 (2016). doi:10.1186/s12931-016-0449-x

42. Taggart, C. C. et al. Elastolytic proteases: inflammation resolution and dysregulation in chronic infective lung disease. Am. J. Respir. Crit. Care Med. 171, 1070–1076 (2005). doi:10.1164/rccm.200407-881PP

43. Pu, S. et al. Effect of sivelestat sodium in patients with acute lung injury or acute respiratory distress syndrome: a meta-analysis of randomized controlled trials. BMC Pulm. Med. 17, 148 (2017). doi:10.1186/s12890-017-0498-z

44. Lu, X. et al. The role of fatty acid metabolism in acute lung injury: a special focus on immunometabolism. Cell. Mol. Life Sci. 81, 120 (2024). doi:10.1007/s00018-024-05131-4

45. Chung, K. P. et al. Alveolar epithelial cells mitigate neutrophilic inflammation in lung injury through regulating mitochondrial fatty acid oxidation. Nat. Commun. 15, 7241 (2024). doi:10.1038/s41467-024-51683-1

46. Williams, A. E. et al. Evidence for chemokine synergy during neutrophil migration in ARDS. Thorax 72, 66–73 (2017). doi:10.1136/thoraxjnl-2016-208597

47. Ward, P. A. The harmful role of C5a on innate immunity in sepsis. J. Innate Immun. 2, 439–445 (2010). doi:10.1159/000317194

48. Bastarache, J. A. et al. Procoagulant alveolar microparticles in the lungs of patients with acute respiratory distress syndrome. Am. J. Physiol. Lung Cell Mol. Physiol. 297, L1035–L1041 (2009). doi:10.1152/ajplung.00214.2009

49. Engelmann, B. & Massberg, S. Thrombosis as an intravascular effector of innate immunity. Nat. Rev. Immunol. 13, 34–45 (2013). doi:10.1038/nri3345

50. Camprubi-Rimblas, M. et al. Anticoagulant therapy in acute respiratory distress syndrome. Ann. Transl. Med. 6, 36 (2018). doi:10.21037/atm.2018.01.08

51. Yang, Z. et al. Complement as a vital nexus of the pathobiological connectome for acute respiratory distress syndrome: an emerging therapeutic target. Front. Immunol. 14, 1100461 (2023). doi:10.3389/fimmu.2023.1100461

52. Camprubi-Rimblas, M. et al. Effects of nebulized antithrombin and heparin on inflammatory and coagulation alterations in an acute lung injury model in rats. J. Thromb. Haemost. 18, 571–583 (2020). doi:10.1111/jth.14685

53. Conway Morris, A., et al. C5a mediates peripheral blood neutrophil dysfunction in critically ill patients. Am. J. Respir. Crit. Care Med. 180, 19–28 (2009). doi:10.1164/rccm.200812-1928OC

54. Poddar, D. et al. An extraribosomal function of ribosomal protein L13a in macrophages resolves inflammation. J. Immunol. 190, 3600–3612 (2013). doi:10.4049/jimmunol.1201933

55. Dai, M.-S. et al. Ribosomal protein L23 activates p53 by inhibiting MDM2 function in response to ribosomal perturbation but not to translation inhibition. Mol. Cell Biol. 24, 7654–7668 (2004). doi:10.1128/MCB.24.17.7654-7668.2004

56. Kruiswijk, F., Labuschagne, C. F. & Vousden, K. H. p53 in survival, death and metabolic health: a lifeguard with a licence to kill. Nat. Rev. Mol. Cell Biol. 16, 393–405 (2015). doi:10.1038/nrm4007

57. Warner, J. R. & McIntosh, K. B. How common are extraribosomal functions of ribosomal proteins? Mol. Cell 34, 3–11 (2009). doi:10.1016/j.molcel.2009.03.006

58. Kalil, A. C. et al. Baricitinib plus Remdesivir for hospitalized adults with Covid-19. N. Engl. J. Med. 384, 795–807 (2021). doi:10.1056/NEJMoa2031994

59. Marconi, V. C. et al. Efficacy and safety of baricitinib for the treatment of hospitalised adults with COVID-19 (COV-BARRIER): a randomised, double-blind, parallel-group, placebo-controlled phase 3 trial. Lancet Respir. Med. 9, 1407–1418 (2021). doi:10.1016/S2213-2600(21)00331-3

60. Ayyat, K. S. et al. Reperfusion inflammatory state of human lungs during cellular ex-vivo perfusion. J. Heart Lung Transplant. 38, S241 (2019). doi:10.1016/j.healun.2019.01.594

61. Yip, S., Wang, N. & Sugimura, R. Give Them Vasculature and Immune Cells: How to Fill the Gap of Organoids. Cells Tissues Organs 212, 369–382 (2023). doi:10.1159/000529431

62. Kollareth, D. J. M. & Sharma, A. K. Precision cut lung slices: an innovative tool for lung transplant research. Front. Immunol. 15, 1504421 (2024). doi:10.3389/fimmu.2024.1504421

63. Granton, E., Kim, J. H., Podstawka, J. & Yipp, B. G. The lung microvasculature is a functional immune niche. Trends Immunol. 39, 890–899 (2018). doi:10.1016/j.it.2018.09.002

64. Zamora, M. E. et al. Marginated neutrophils in the lungs effectively compete for nanoparticles targeted to the endothelium, serving as a part of the reticuloendothelial system. ACS Nano 18, 22275–22297 (2024). doi:10.1021/acsnano.4c06286

65. Craven, T. H. et al. Activated neutrophil fluorescent imaging technique for human lungs. Sci. Rep. 11, 976 (2021). doi:10.1038/s41598-020-80083-w

66. Humphries, D. C. et al. Specific in situ immuno-imaging of pulmonary-resident memory lymphocytes in human lungs. Front. Immunol. 14, 1100161 (2023). doi:10.3389/fimmu.2023.1100161

67. Thompson, B. T., Chambers, R. C. & Liu, K. D. Acute respiratory distress syndrome. N. Engl. J. Med. 377, 562–572 (2017). doi:10.1056/NEJMra1608077

