## Supplemental fig 1 for "Therapeutic signature mapping of paired direct and indirect LPS injury in an ex vivo human lung perfusion platform reveals injury-specific druggable programs"

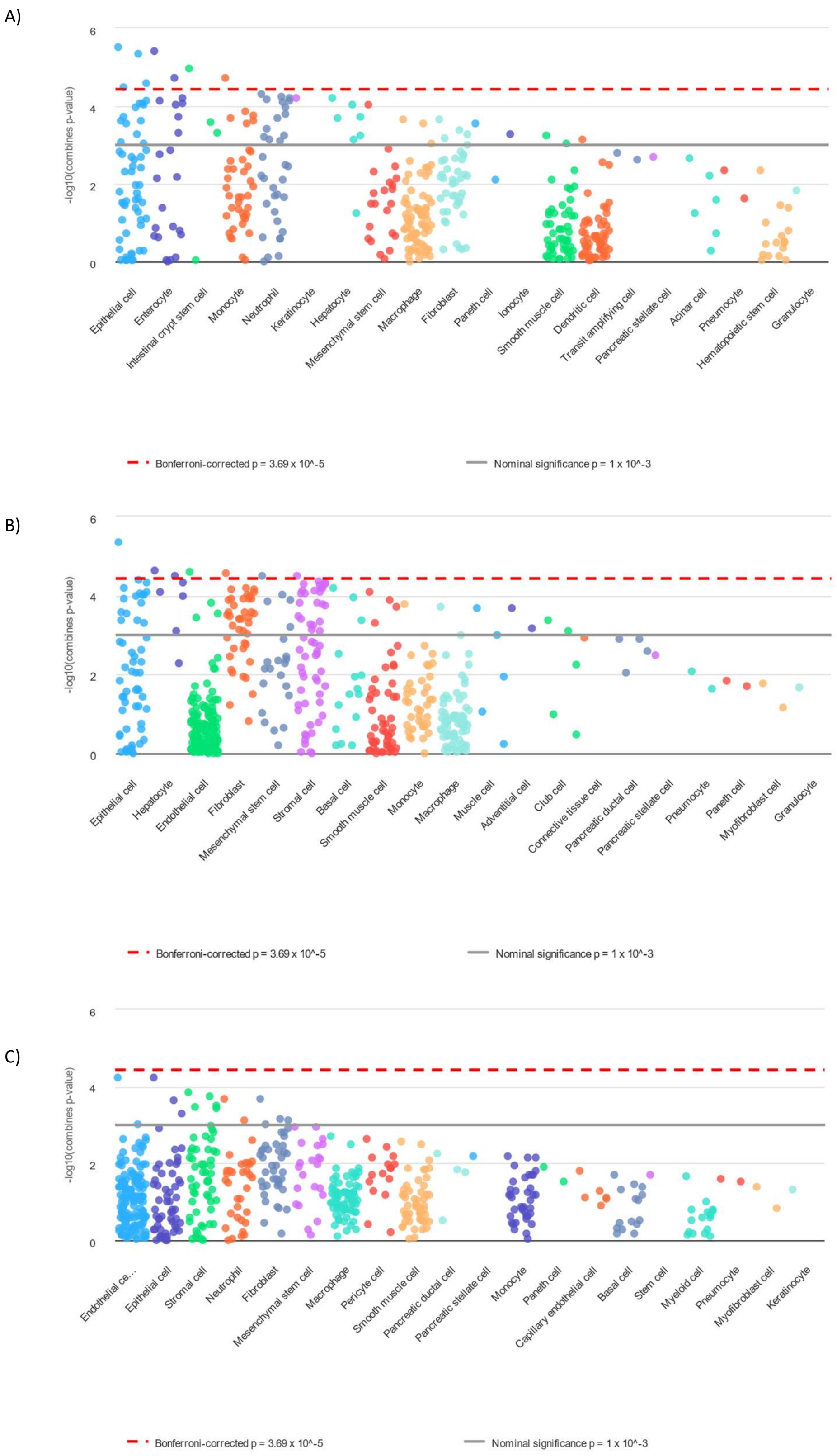


**Supplementary Figure 1: Cell-type enrichment using WebSCEA**: Manhattan plot of the top 20 cell types over enriched for differentially identified proteins in: A) Direct injury. B) Indirect injury. C) EVLP comparison
