## Supplemental fig 2 for "Therapeutic signature mapping of paired direct and indirect LPS injury in an ex vivo human lung perfusion platform reveals injury-specific druggable programs"

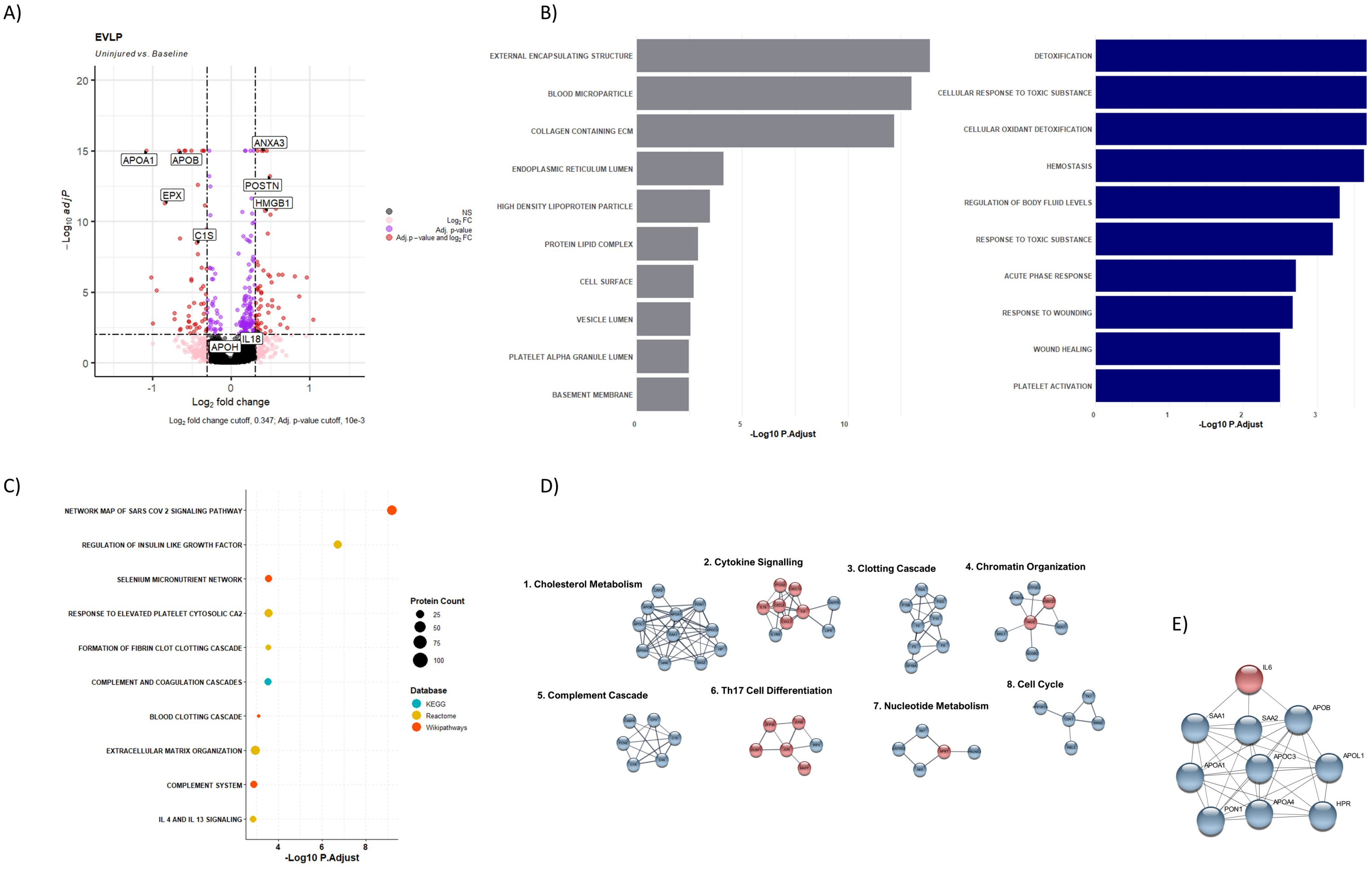


**Supplementary figure 2: Effect of EVLP:** A) Volcano plot of EVLP comparison between uninjured group vs baseline group. X-axis represents the log2 foldchange while the y-axis the -log10 q value. Circles in red were significant based on cut-off while pink had changed in log2 foldchange but not q-value while blue had a significant q-value but no change in the log2foldchange. B) Bar chart of EVLP proteins functional annotation using GO, grouped by category grey representing cellular component and blue biological process. X-axis -log10 adjusted p value for the pathways tested. C) Dot plot of EVLP proteins pathway analysis results, color of dot represents source database and size represents number of proteins within that pathway. X-axis -log10 adjusted p value for the pathways tested. D) Functional clusters within EVLP PPI network with ≥5 proteins. E) EVLP PPI network of the hub genes identified.
