## Supplemental table 1 for "Therapeutic signature mapping of paired direct and indirect LPS injury in an ex vivo human lung perfusion platform reveals injury-specific druggable programs"

| Sex | Age | Donation after circulatory death | Cause of Death | Smoker | Height | Weight | Cold ischaemic time (hrs) | Respiratory failure severity |
|---|---|---|---|---|---|---|---|---|
| M | 55 | No | Intracranial Haemorrhage | Yes | 179 | 82 | 24 | Moderate |
| M | 23 | Yes | Anoxia | No | 183 | 75 | 72 | Mild |
| M | 58 | Yes | Anoxia | No | 170 | 92.6 | 108 | Mild |
| M | 27 | No | Head Trauma | No | 180 | 90 | 72 | Mild |
| M | 59 | Yes | Stroke | No | 168 | 93 | 60 | Mild |
| F | 22 | No | Anoxia | Yes | 160 | 59 | 96 | None |
| F | 70 | No | Intracranial Haemorrhage | No | 165 | 100 | 60 | None |
| M | 32 | No | Anoxia | No | 162 | 82 | 72 | None |
| F | 62 | Yes | Intracranial Haemorrhage | No | 166 | 85 | 18 | None |
| F | 48 | No | Intracranial Haemorrhage | Yes | 174 | 85 | 60 | Moderate |

**Supplementary Table 1**: Clinical details of donors.
