## Supplemental table 2 for "Therapeutic signature mapping of paired direct and indirect LPS injury in an ex vivo human lung perfusion platform reveals injury-specific druggable programs"

Supplementary Table 2 | Therapeutic signature reversal highlights injury-specific candidate drug classes

Summary of CLUE L1000-derived signature reversal results stratified by injury model. Norm_cs, normalised connectivity score (more negative indicates stronger predicted reversal of the query signature in the cell line).

| Injury model | Signature layer | Signature direction | Top predicted reversal compounds (cell line; norm_cs) | Convergent mechanistic theme |
|---|---|---|---|---|
| Direct | Proteomics | Upregulated proteins | WT-171 (HCC515; −2.02); Scriptaid (A549; −1.98) | HDAC inhibition — the only lung-specific reversal signal for direct epithelial stress |
| Direct | Proteomics | Downregulated proteins | GNF-2 (NPC; −1.72); ABT-737 (MCF7; −1.67); Rosiglitazone (NPC; −1.66) | BCL/survival and PPARγ/metabolic restoration |
| Indirect | Proteomics | Upregulated proteins | Mebendazole (HELA; −2.31); Olmesartan (HELA; −2.25); Flubendazole (A549; −2.14, lung); CYT-387 (A549; −2.00, lung) | Tubulin/cytoskeletal and RAAS reversal overall; JAK/STAT inhibition lung-specific |
| Indirect | Proteomics | Downregulated proteins | Ruxolitinib (A375; −1.64); MLN-0128 (THP1; −1.62); GSK-690693 (PC3; −1.59). Lung: Pterostilbene (A549; −1.56); Prednisolone (HCC515; −1.52) | JAK/STAT and mTOR/AKT pathway reversal; lung-specific glucocorticoid and metabolic restoration |
