## Supplemental table 3 for "Therapeutic signature mapping of paired direct and indirect LPS injury in an ex vivo human lung perfusion platform reveals injury-specific druggable programs"

### Supplementary Table 3 | Cross-platform class concordance summary

Platform-level concordance across CLUE L1000, Enrichr, L1000CDS² and Open Targets for major therapeutic classes nominated by EVLP-derived signatures.

| Drug/target class | CLUE | Enrichr | L1000CDS² | Open Targets |
|---|---|---|---|---|
| JAK/STAT inhibitors | lung-specific hit(s) reported | validated | present | strong respiratory associations |
| PI3K/AKT/mTOR inhibitors | top-ranked kinase hits | mTOR/PI3K hits | dominant | ALI/respiratory evidence |
| SYK inhibitors | top-ranked (direct) | present | top-ranked (direct) | pneumonia association |
| CDK inhibitors | not present | strong enrichment | validated | tractable target evidence |
| HDAC inhibitors | lung-specific reversal (direct UP) | validated | not top-ranked | supportive/limited |
| PPARγ agonists | validated | dominant | not top-ranked | metabolic context |
